# Challenges and advances for huntingtin detection in cerebrospinal fluid: in support of relative quantification

**DOI:** 10.1101/2024.09.25.614766

**Authors:** Rachel J. Harding, Yuanyun Xie, Nicholas S. Caron, Hailey Findlay-Black, Caroline Lyu, Nalini Potluri, Renu Chandrasekaran, Michael R. Hayden, Blair R. Leavitt, Douglas R. Langbehn, Amber L. Southwell

## Abstract

Huntington disease (HD) is a progressive and devastating neurodegenerative disease caused by expansion of a glutamine-coding CAG tract in the huntingtin (*HTT*) gene above a critical threshold of ∼35 repeats resulting in expression of mutant HTT (mHTT). A promising treatment approach being tested in clinical trials is HTT lowering, which aims to reduce levels of the mHTT protein. Target engagement of these therapies in the brain are inferred using antibody-based assays to measure mHTT levels in the cerebrospinal fluid (CSF), which is frequently reported as absolute mHTT concentration based on a monomeric protein standard used to generate a standard curve. However, patient biofluids are a complex milieu of different mHTT protein species, suggesting that absolute quantitation is challenging, and a single, recombinant protein standard may not be sufficient to interpret assay signal as molar mHTT concentration. In this study, we used immunoprecipitation and flow cytometry (IP-FCM) to investigate different factors that influence mHTT detection assay signal. Our results show that HTT protein fragmentation, protein-protein interactions, affinity tag positioning, oligomerization and polyglutamine tract length affect assay signal intensity, indicating that absolute HTT quantitation in heterogeneous biological samples is not possible with current technologies using a single standard protein. We also explore the binding specificity of the MW1 anti-polyglutamine antibody, commonly used in these assays as a mHTT-selective reagent and demonstrate that mHTT binding is preferred but not specific. Furthermore, we find that MW1 depletion is not only incomplete, leaving residual mHTT, but also non-specific, resulting in pull down of some wildtype HTT protein. Based on these observations, we recommend that mHTT detection assays report only relative mHTT quantitation using normalized arbitrary units of assay signal intensity, rather than molar concentrations, in the assessment of central nervous system HTT lowering in ongoing clinical and preclinical studies, and that MW1-depletion not be used a method for quantifying wildtype HTT protein.

## Introduction

Huntington disease (HD) is a devastating, inherited neurodegenerative disease with progressive cognitive, psychological and physical symptoms. HD is caused by expansion of the CAG repeat tract in exon 1 of the huntingtin (*HTT*) gene above a critical threshold of ∼35 repeats, resulting in expression of a polyglutamine (polyQ) expanded form of the HTT protein, referred to as mutant HTT (mHTT)(The Huntington’s Disease Collaborative Research Group, 1993). HTT plays important roles in proteostasis (Harding and Tong, 2018), axonal transport (Vitet et al., 2020), transcription regulation (Benn et al., 2008), cellular stress responses (Liu and Zeitlin, 2017), and mitochondrial function (Carmo et al., 2018) and the expression of mHTT is considered responsible for the molecular pathogenesis cascade, including both loss of function and gain of toxic function, resulting in HD phenotypes in HD animal models and patients. However, despite being a monogenic disorder, the mechanisms of HD pathophysiology are complex, and remain the subject of intense study (Saudou and Humbert, 2016).

The majority of candidate therapies currently being tested in clinical trials for HD aim to lower levels of the mHTT protein (Tabrizi et al., 2019). To infer target-engagement of these drugs, mHTT levels, usually from the cerebrospinal fluid (CSF), are monitored using ultrasensitive detection assays (Southwell et al., 2015; Wild et al., 2015). Decreased mHTT levels in HD model mouse brain following intracerebroventricular administration of *HTT*-targeting antisense oligonucleotides (ASOs) were shown to induce correlative mHTT lowering in CSF, validating CSF mHTT quantitation as a pharmacodynamic biomarker for HTT lowering clinical trials (Southwell et al., 2015). mHTT is also a monitoring biomarker for HD, and its levels track with proximity to disease onset as well as cognitive and motor deficits (Southwell et al., 2015; Wild et al., 2015). Numerous different mHTT detection assays have been developed to date (Fodale et al., 2017; Landles et al., 2021; Reindl et al., 2019; Southwell et al., 2015; Weiss et al., 2009; Wild et al., 2015), all of which employ capture-probe antibody pairs, one which is used to immunoprecipitate and the other is used for detection of mHTT for biofluid samples. Moreover, a single full-length mHTT protein standard is often used to determine a molar concentration of mHTT from assay signal.

mHTT exists in many different proteoforms including alternatively spliced fragments (Neueder et al., 2017), proteolytically cleaved fragments (El-Daher et al., 2015; Graham et al., 2006; Landles et al., 2010), in complexes with a myriad of binding partners (Greco et al., 2022; Harding et al., 2021; Ratovitski et al., 2012) and with different polyQ tract lengths generated via somatic expansion mechanisms (Aviolat et al., 2019; Telenius et al., 1994). However, our current understanding of the relative distribution of these proteoforms in biofluids and other samples from people with HD or HD animal models, or how they track with disease, remains limited.

One limitation of mHTT detection immunoassays is the inherent bias in which specific proteoforms are detected, which is defined by the epitopes of the antibody pair that are used. Different antibody pairs will preferentially detect different proteoforms of mHTT (Landles et al., 2021) and no one pair of antibodies can detect all or “total” HTT in the complex milieu of species that exists in biological samples given the fragmentation and variety of conformations of this protein. Additionally, the commonly used MW1 antibody, which was raised against the DRPLA-19Q/GST fusion protein, binds polyQ expanded proteins, and can form stoichiometrically heterogenous interactions with mHTT species (Bravo-Arredondo et al., 2023; Owens et al., 2015). This suggests that polyQ length functions as an additional variable for detection assay signal, in addition to mHTT concentration. Indeed, the variable stoichiometry of MW1-mHTT polyQ interactions compared to other anti-HTT antibodies likely accounts for the so-called detection paradox where mHTT concentration exceeds total HTT concentration (Fodale et al., 2020). This paradox suggests that absolute quantitation of mHTT may not be possible with immunoassays with a single protein standard across an array of patient samples where CAG number, and hence polyQ length, is variable, and that defining molar concentrations of mHTT by such a methodology may be misleading.

In this study, we set out to explore different factors which can influence mHTT detection assay signal using a suite of HTT proteins, including a comprehensive allelic series of full-length HTT samples, spanning wildtype to juvenile HD polyQ tract lengths. Employing an IP-FCM assay, as well as other immunoassay approaches, we show that a variety of mHTT properties and assay condition considerations influence assay signal and show that using a single protein standard across an array of biological samples is not sufficient to allow accurate calculation of mHTT concentration. We also explore the binding specificity of the MW1 antibody and demonstrate that detection of polyQ expanded mHTT is preferential but not specific. Together these data support a new paradigm for mHTT detection, where results are reported as relative quantitation in reference to a given standard protein rather than reporting absolute concentrations. Moreover, that MW1-depletion should not be used to quantify wildtype HTT protein.

## Results

### Purification of an allelic series of full-length HTT protein samples

Previously, we designed and developed an open-source toolkit for the eukaryotic expression and purification of full-length HTT proteins with different polyQ tract lengths and N-or C-terminal FLAG tags for purification and/or detection (Harding et al., 2019). This toolkit is a unique resource for HD research as it encompasses a fine-grain allelic series of HTT proteins corresponding to wildtype control (Q23, Q25, Q30), HD threshold inflection point (Q36), adult-onset HD (Q42, Q52, Q54) and juvenile-onset HD (Q60, Q66). Regardless of polyQ tract length, all proteins can be co-expressed with HAP40, an important interaction partner of HTT whose levels track with HTT in cells and which functions to stabilize the large HTT protein molecule (Harding et al., 2021; Huang et al., 2021a, 2021b; Xu et al., 2022).

All HTT proteins were purified from insect cells using a two-step protocol, FLAG-affinity chromatography and gel filtration, and verified by SDS-PAGE (**Supplementary Figure 1**). We have previously validated HTT and HTT-HAP40 samples produced using this toolkit with numerous biophysical and structural methodologies (Harding et al., 2021, 2019) to show they are pure, monodisperse, folded and functional samples amenable to downstream interrogation. We further complemented this suite of HTT proteins with mHTT fragment proteins described previously (Southwell et al., 2015), to ensure better coverage of the milieu of mHTT species which are present in patient and HD animal model biofluid samples. The fragment proteins included a construct spanning aa. 1-171 with Q68 fused to aa. 1744-2234, hereafter called fusion HTT Q68, and aa. 1-586 with Q68, hereafter called N586 HTT Q68.

**Figure 1.**
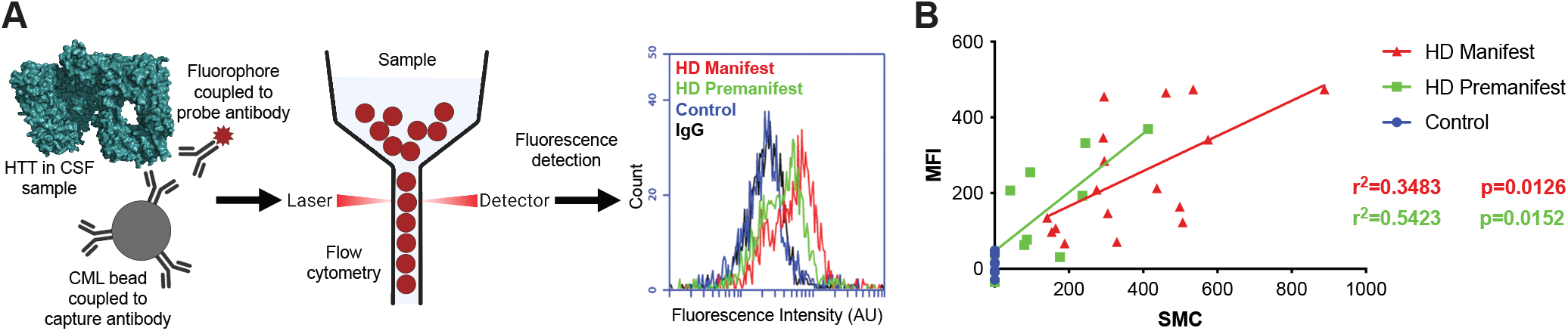
IP-FCM to detect mHTT and comparison of assay with Singulex approach. A. IP-FCM assay workflow to detect mHTT in biofluid samples. B. Head-to-head comparison of IP-FCM with Singulex mHTT detection assay approaches with the same biofluid sample set from control, premanifest, and manifest HD participants shows good agreement of the methodologies. R-squared and p-values calculated from simple linear regression analysis.

### HDB4/MW1 IP-FCM assay for ultrasensitive detection of HTT shows that protein concentration and polyQ tract length influence assay signal

To measure the signal elicited from our panel of HTT proteins under different conditions, we used a micro-bead-based immunoprecipitation-flow cytometry (IP-FCM) assay. This assay was previously optimized, and a capture-probe antibody pair were identified which permit ultrasensitive detection of mHTT in HD mouse model and patient CSF (**Figure 1A**)(Southwell et al., 2015). Our IP-FCM assay employs the HDB4E10 antibody (hereafter HDB4), which was raised against an epitope within the bridge domain aa. 1844-2131 of HTT and also recognizes an epitope in exon 1 (**Supplementary Figure 2**), and the polyQ-specific MW1 antibody which is employed in other published mHTT detection assays (Fodale et al., 2017; Landles et al., 2021; Reindl et al., 2019; Weiss et al., 2009; Wild et al., 2015). Assay signal from our HDB4/MW1 IP-FCM assay and a different ultrasensitive mHTT detection assay also suitable for use in CSF that employs Singulex technology, showed statistically significant correlation of mHTT assay signal (**Figure 1B**).

**Figure 2.**
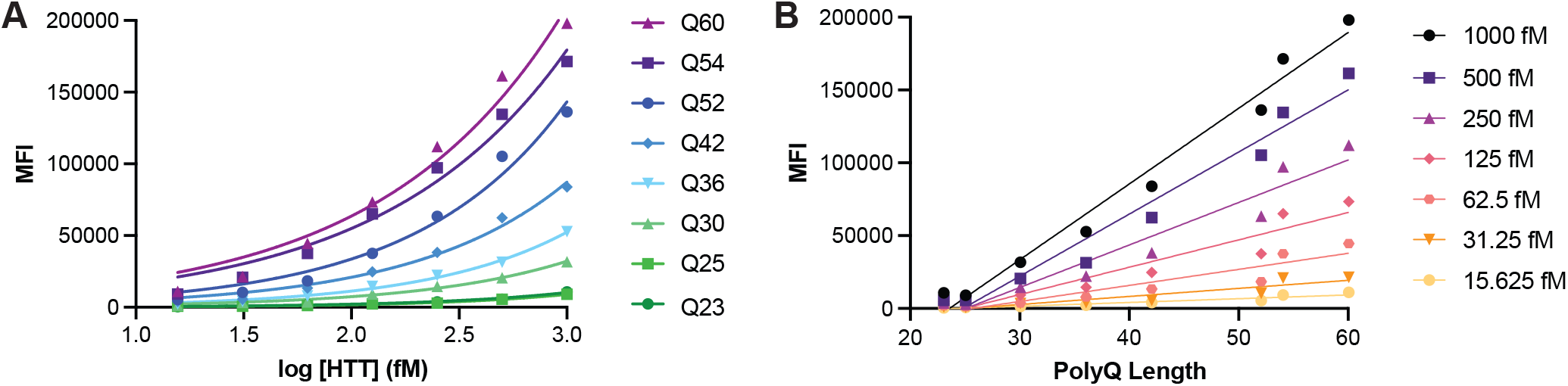
HTT protein concentration and polyQ tract length influence IP-FCM assay signal. HDB4/MW1 IP-FCM assay analysis of C-terminal FLAG tagged full-length HTT with polyQ tract lengths spanning Q23 to Q60. Assay signal (mean fluorescence intensity – MFI) is plotted as A. a function of protein concentration or B. polyQ tract length. Graphs shown are generated from a representative replicate dataset, N=3.

Analysis of a titration of our allelic series of purified full-length HTT samples using the IP-FCM assay reveals that assay signal is modulated by polyQ length as well as concentration, with greater assay signal measured for higher protein concentrations and longer polyQ tract lengths (**Figure 2A**). Replotting these data as a function of polyQ tract length shows that the relationship between polyQ length and assay signal at a defined protein concentration is approximately linear under the conditions tested (**Figure 2B**).

### Different structural properties of the HTT protein can influence IP-FCM assay signal

Next, we investigated how different structural features of the HTT protein might influence HDB4/MW1 IP-FCM assay signal. Beyond polyQ tract length, HTT proteoform heterogeneity in HD patient CSF is expected due to alternatively spliced fragments (Neueder et al., 2017), or fragmentation due to proteolytic cleavage (El-Daher et al., 2015; Graham et al., 2006; Landles et al., 2010), as well as aggregation of the protein into higher order oligomers (Tan et al., 2015). Recombinant proteins used as standards can also differ by the position of their purification tags which can further modulate their structure and/or conformation. Using a suite of HTT proteins, we investigated these variables.

Firstly, full-length and fragment HTT proteins bearing approximately the same polyQ tract were assessed using the HDB4/MW1 IP-FCM assay, revealing significantly different profiles for each protein over equivalent concentration titrations (**Figure 3A**). Despite all three proteins containing approximately the same polyQ-tract length, the assay signal is influenced by the context of these epitopes with the greatest signal observed in the full-length protein, potentially due to conformational flexibility and/or other structural changes that different epitope flanking sequences confer to the protein molecule.

**Figure 3.**
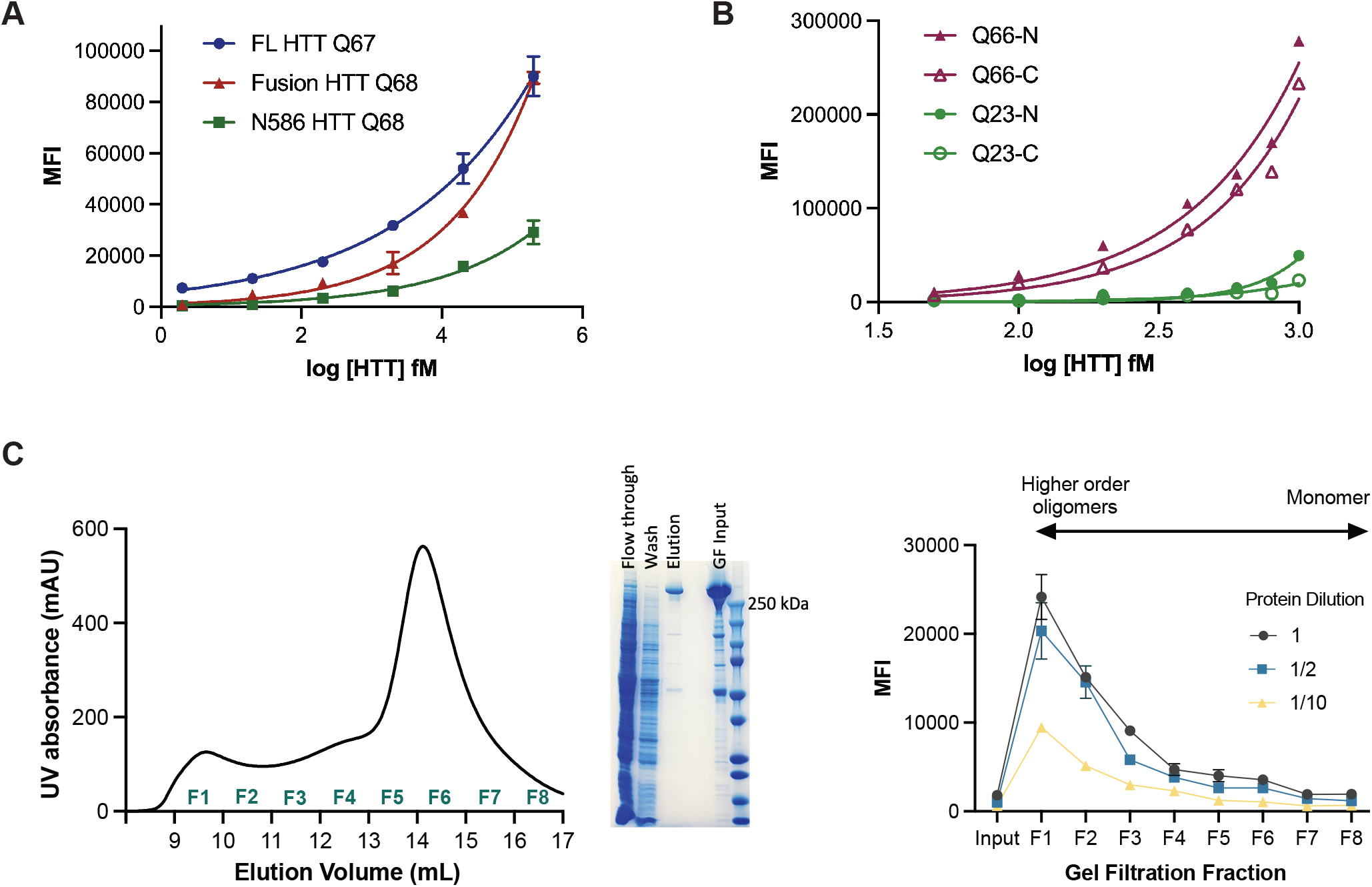
IP-FCM HTT detection assay signal is influenced by protein fragmentation, the position of the affinity tag and the oligomerization state of the protein. A. MW1-HDB4 IP-FCM analysis of full-length (FL) HTT, fusion HTT Q68 and N586 HTT Q68 protein with approximately the same Q-length. B. HDB4/MW1 IP-FCM analysis of full-length HTT with polyQ tracts spanning either 23 or 66 glutamines, with N or C-terminal FLAG-tag. C. Left – Gel filtration (GF) trace of FLAG-affinity chromatography purified full-length HTT Q54 applied to Superose6 10/300 GL column which elutes across fractions 1-8 (F1-F8). Middle – SDS-PAGE analysis of FLAG-affinity chromatography flow through, wash, elution and GF input fractions. Right - HDB4/MW1 IP-FCM analysis of concentration normalized GF fractions F1-F8 at different dilutions. IP-FCM graphs shown are generated from a representative replicate dataset, N=3.

Next, we assessed full-length HTT proteins, with either N-or C-terminal FLAG tags and polyQ tract lengths of Q23 or Q66 using the HDB4/MW1 IP-FCM assay (**Figure 3B**). For HTT Q66, proteins bearing a C-terminal FLAG tag compared to an N-terminal FLAG tag show lower assay signal. A similar trend is also seen for HTT Q23 at higher concentrations of protein, with N-terminally tagged HTT Q23 eliciting higher assay signal that C-terminally tagged HTT Q23. This finding suggests that affinity tag positioning can alter the accessibility or conformation of antibody epitopes. Indeed, the MW1 epitope begins just 17 residues after the N-terminal tag and flanking sequence composition is known to influence the biophysical properties of the polyQ tract (Duennwald et al., 2006; Shen et al., 2016).

We then looked at the effects of HTT protein oligomerization and aggregation on IP-FCM assay signal. The gel filtration elution profile of apo HTT has a distinct shape, reported by multiple groups (Harding et al., 2019; Huang et al., 2015; Kim et al., 2021; Pace et al., 2021). The main peak corresponding to monomeric HTT eluting after ∼0.6 column volumes preceded by dimer, tetramer and increasingly higher order oligomer peaks eluting ahead of the monomer peak, with the largest oligomers and aggregates eluting in the column void volume (∼0.3 column volumes). FLAG-affinity chromatography purified HTT Q54 (∼85% pure) was concentrated (input) (**Figure 3C**, middle) and applied to Superose6 Increase 10/300 column, then we collected eight 1 mL fractions spanning all peaks (fractions 1-8) (**Figure 3C**, left). The concentration of the eight gel filtration fractions and input sample was normalised and then samples of each analysed by HDB4/MW1 IP-FCM assay at different dilutions (**Figure 3C**, right). Fractions corresponding to mHTT monomer yielded the lowest signal in this assay (F6-8), with signal increasing as oligomeric state increased to the largest assemblies (F1). The change in assay signal with oligomeric state might reflect avidity effects occurring in higher order assemblies of HTT protein, where adjacent epitopes across protein molecules are more likely to be in closer proximity, as is reported for other aggregate protein immunoassays (Pan et al., 2005).

### IP-FCM assay buffer can influence assay signal for some HTT protein complexes

HTT is a protein scaffold and is reported to bind more than 500 proteins (Greco et al., 2022). Because of the varying interaction interfaces and conformational changes induced by complex formation, we hypothesised that different protein complexes of HTT are likely to have altered epitope availability. HAP40 is the only structurally validated interaction partner of HTT and can bind HTT with either wildtype (Q23) or disease expanded (Q54) polyQ tract lengths (Harding et al., 2021; Huang et al., 2021a). Assay buffer components, such as detergents, can influence protein complex structure and stability. Performing our IP-FCM assay with a buffer more closely resembling physiological conditions, such as artificial cerebrospinal fluid (aCSF) (**Figure 4A**), a difference in assay signal can be observed for apo compared to HAP40-bound forms of HTT for the Q54 form of the protein (p<0.0001 for all concentrations, 2way ANOVA multiple comparisons of log-log data) but not Q23, indicating that expanded exon 1 might be differently structured in these two proteoforms of HTT. However, when the assay is performed under more stringent buffer conditions, such as with the detergent NP-40 (1% (v/v/)) (**Figure 4B**), this difference in signal is lost. This could be due to NP-40-mediated disruption of the HTT-HAP40 complex itself or conformational uniformity of exon 1 structure for apo and HAP40-bound HTT in the presence of detergent.

**Figure 4.**
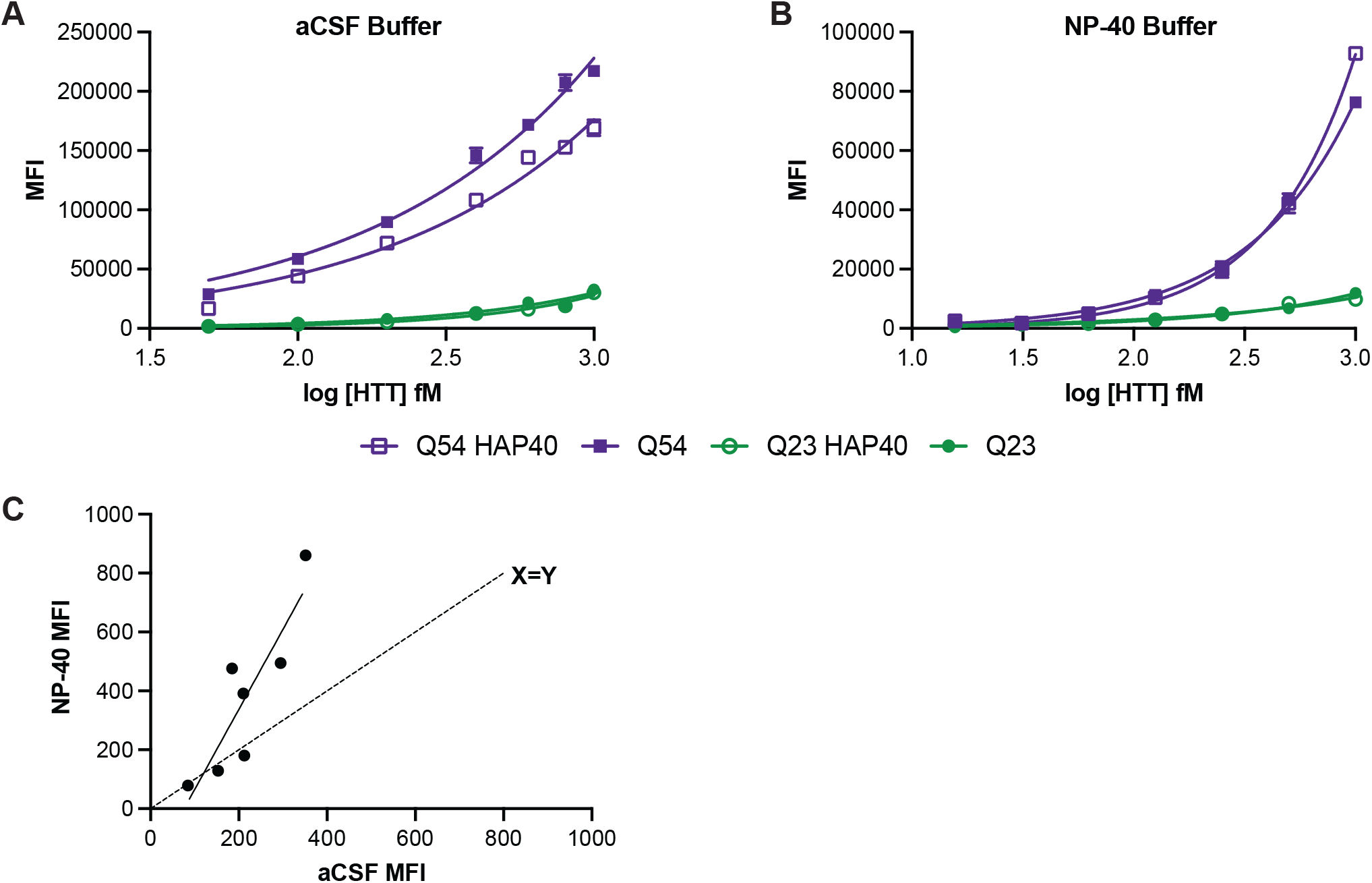
Detergent can alter IP-FCM assay signal for some HTT proteins. HDB4/MW1 IP-FCM analysis of HTT and HTT-HAP40 with either Q23 or Q54 in A. artificial CSF (aCSF) or B. 1% (v/v) NP-40-containing buffer. Graphs shown are generated from a representative replicate dataset, N=3. C. Comparison of assay signal obtained from human CSF samples diluted 1:1 in either aCSF and or NP-40 buffer.

Next, we compared the assay signal obtained from human CSF samples diluted 1:1 in either aCSF or a buffer containing NP-40 detergent (**Figure 4C**). Although samples with relatively low assay signal gave comparable signal in both buffer conditions, as shown by their proximity to the X=Y line, those with higher mHTT concentration as denoted by higher assay signal showed much higher signal in NP-40 buffer compared to aCSF. Again, this indicates that detergent in assay buffer can modulate epitope availability and binding by the HDB4/MW1 antibody pair in the IP-FCM assay format, perhaps related to changes in HTT complex formation, conformation or structure under different buffer conditions.

### MW1 has preference but not specificity for mHTT and can deplete wildtype HTT

The MW1 antibody is also used to pretreat samples for analysis in ultrasensitive wildtype HTT detection assays, with the aim of depleting biofluid samples of mHTT, leaving only wildtype HTT to be detected with polyQ length-independent immunoassay antibody pair (Boyanapalli et al., 2022).

However, our HTT allelic series IP-FCM data (**Figure 2**) show that MW1 has only preference, not specificity for mHTT, suggesting that such an assay approach may also deplete wildtype HTT. This corroborates previously studies which drew the same conclusion (Bennett et al., 2002; Owens et al., 2015).

We conducted a parallel experiment with Hu97/18 mouse CSF depleted using MW1 and assessed the depleted CSF and MW1 IP fraction using HDB4/MW1 IP-FCM. The depleted CSF shows reduced signal in the assay compared to the sample obtained by MW1 IP (**Figure 5B**). This finding confirms that MW1 binds both HTT Q18 and Q97 in this experiment, and binding preference, but not complete specificity is shown by MW1 to mHTT. The degree to which each form of HTT is depleted from a biofluid samples likely depends on the relative affinity of MW1 for the specific polyQ tract lengths of the HTT proteins in question, as well as avidity effects, which will be driven by the relative concentrations of the proteins and the MW1 antibody in the depletion experiment conditions.

**Figure 5.**
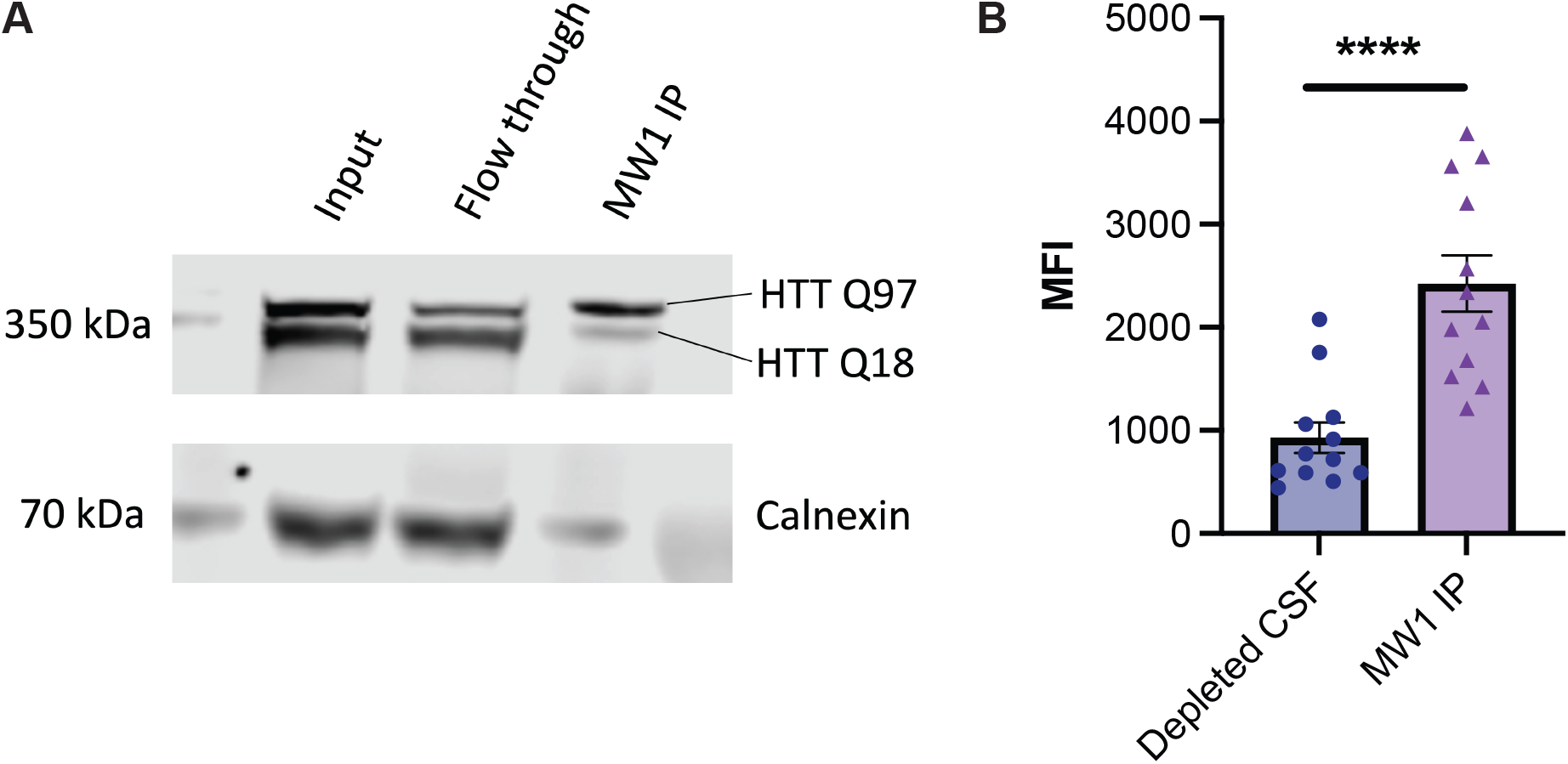
MW1 depletion of mHTT is both incomplete and non-specific in HD mouse brain lysate and CSF. A. HTT allele separation Western blot analysis of Hu97/18 brain lysate used for MW1 immunoprecipitation (IP). Fractions corresponding to the IP input, flow through (depleted) and elution (IP captured) are shown, with Calnexin as a control. B. HDB4/MW1 IP-FCM analysis of Hu97/18 mouse CSF after depletion by MW1 shows residual mHTT protein in depleted CSF.

To test this hypothesis, we first used MW1 in an immunoprecipitation (IP) experiment with Hu97/18 HD model mouse brain lysates. Hu97/18 mice express full-length human HTT Q97 and Q18 and lack mouse *Hdh* (**Figure 5A**) (Southwell et al., 2013). Analysis of the input, flow through and IP fraction using a western blot that separates wildtype and mHTT bands (Carroll et al., 2011), shows that the mHTT protein is diminished in the flow through fraction compared to input, though not completely depleted, and that both forms of the protein are present in the elution.

### The affinity of MW1 for HTT is influenced by polyQ length

To further investigate MW1 binding to our full-length HTT allelic series, we used two orthogonal assays to measure MW1 interaction with different polyQ length HTT proteins. MW1-HTT binding was analysed under native and denatured conditions using enzyme-linked immunosorbent assay (ELISA) and western blot analysis respectively. For ELISA, HTT was adhered to the plate surface and a titration of MW1 antibody was incubated prior to detection with HRP-linked secondary antibody as previously described (Denis et al., 2023). The MW1 titration was optimised to ensure binding saturation as seen by stabilised A450 readings as MW1 concentration increases (**Figure 6A**). This permitted calculation of binding affinities of MW1 for each HTT protein, reported as apparent KD values (Kapp)(**Figure 6B**), as the stoichiometry of binding complex for each polyQ tract length is unknown and varies as a function of the levels of HTT immobilised on the plate surface. In this assay, A450 values change as a function of MW1 concentration and polyQ tract length. This finding parallels our earlier observations of small Q-length changes influencing immunoassay signal in our IP-FCM analysis (**Figure 2**). We observe that Kapp decreases exponentially with polyQ tract length indicating increasingly high affinity binding to longer polyQ tract length HTT proteins. This indicates that, in solution, MW1 binds HTT in a polyQ tract length dependent manner, as others have shown before (Bravo-Arredondo et al., 2023; Li et al., 2007; Owens et al., 2015), but also highlights how MW1 only has preference for mHTT and is not specific for HTT species with polyQ tracts above the disease threshold length.

**Figure 6.**
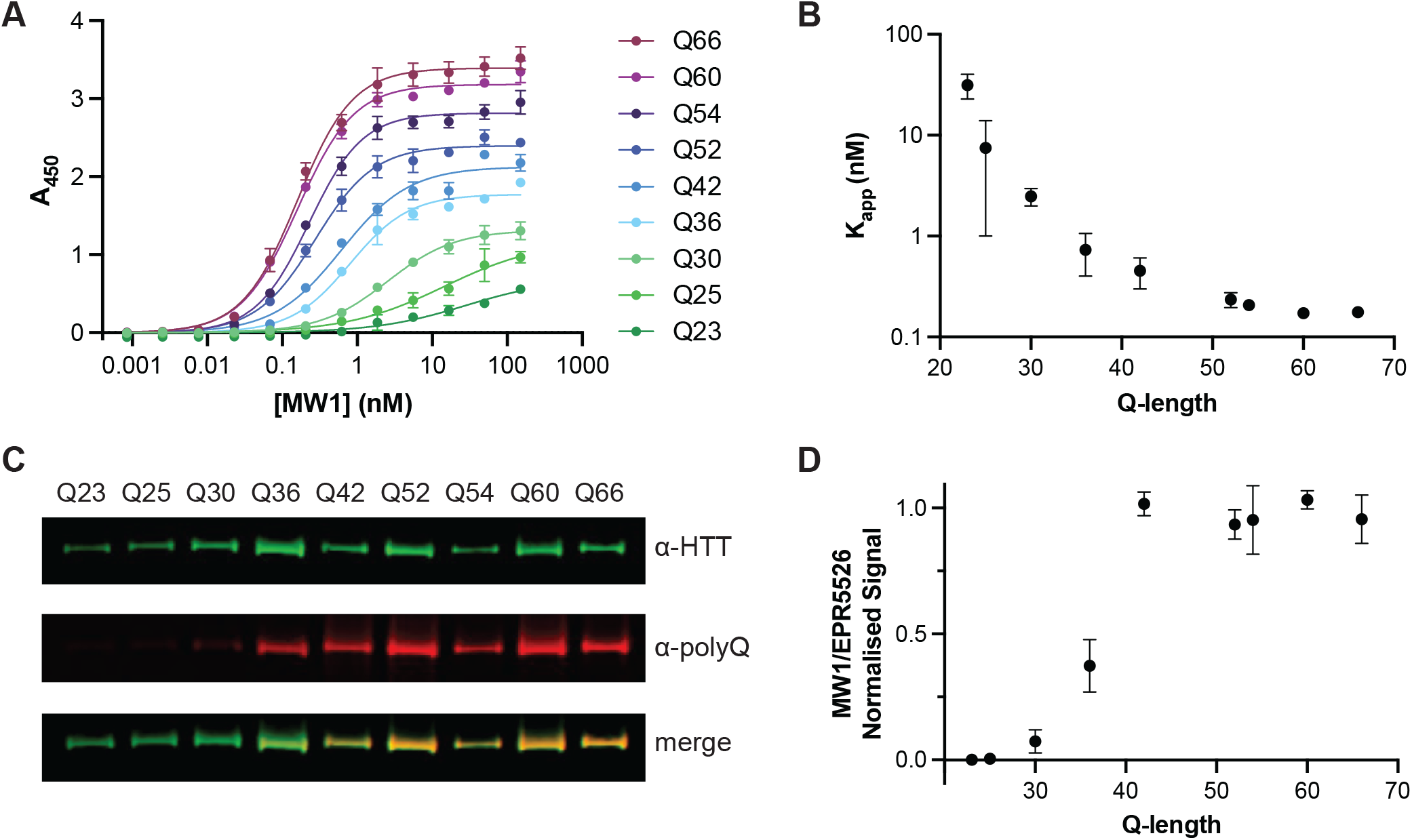
MW1 binding to HTT in different assay formats is dependent on polyQ tract length but is not specific for mHTT. A. Representative ELISA showing binding profile of MW1 to full-length HTT allelic series spanning Q23 to Q66. Error bars are S.D. of three intra-assay replicates. Data fitted in GraphPad Prism with specific binding with hill slope model. B. Mean Kapp (apparent KD) from three independent ELISA replicates plotted as a function of HTT polyQ tract length. Error bars are S.D. of three inter-assay replicates. C. Representative western blot analysis of full-length HTT allelic series spanning Q23 to Q66 with ∼5 ng loaded per lane. Blots probed with both α-HTT EPR5526 and α-polyQ MW1 shown separately and merged. Full data in **Supplementary Figure 4**. D. Mean normalised MW1/EPR5526 signal from three independent western blot replicates plotted as a function of Q-length. Error bars are S.D. of three inter-assay replicates.

To understand if this finding for MW1-epitope interaction is specific to HTT, we next investigated another polyQ tract containing protein, ataxin-3. Using two different polyQ length ataxin-3 proteins, we repeated the ELISA protocol with ataxin-3 Q10 or Q80 adhered to the plate this time. We observe the same pattern for these two proteins with A450 maximal signal greatly increased for Q80 vs Q10 and the calculated Kapp values much lower for Q80 than Q10, indicating much tighter binding (**Supplementary Figure 3**).

In our western blot analysis of the full-length HTT allelic series, approximately equal amounts of each HTT protein were analysed by western blot, probing with MW1 and also EPR5526, a polyQ independent antibody which targets another region of the exon 1 sequence. Similar to our ELISA experiments, we observed polyQ tract length dependent binding of MW1 (**Figure 6C**) with only faint bands observed for HTT proteins with wildtype polyQ tract lengths. Calculating the normalised signal ratio of MW1/EPR5526, we observe an inflection point ∼Q36 with stabilised signal ratio for HTT proteins with polyQ tract >42. Together, this further validates our conclusion that MW1 interaction with HTT is polyQ length driven but not specific for mHTT (**Figure 6D**).

## Discussion

In this study we show that mHTT ultrasensitive detection assay signal is dependent on many factors beyond protein concentration, including fragmentation, protein-protein interaction, affinity tag positioning, oligomerization and polyglutamine tract length. Additionally, we demonstrate that MW1 has preference but not specificity for mHTT and can bind wildtype HTT, albeit with reduced affinity compared to disease-range polyQ tract length proteoforms of mHTT.

### Box 1.

**Recommendations for best practices for reporting ultrasensitive HTT detection assay data**

Many factors influence assay signal when measuring mutant HTT including polyQ tract length, epitope context, oligomerization state and protein-protein interactions so absolute mHTT quantitation in heterogeneous biological samples is not possible with current technologies.

⇒ We recommend that investigators should report relative signal for a given assay run for a defined standard protein, not protein concentration. MW1 depletion of biosamples to measure wildtype HTT are incomplete and preferential rather than specific and likely still measure a mixture of wildtype and mutant HTT species.
⇒ The community needs to consider alternative approaches for reliably measuring wildtype HTT levels. Detergent influences assay signal in HTT detection assays, calling into question common practices of detergent addition prior to long term storage, which would impact the ability to test-retest samples
⇒ Detergent should be consistently added to samples prior to testing and long-term storage

Biological fluid samples from HD animal models and patients, such as CSF, contain a heterogenous mix of mHTT and wildtype HTT species, including fragments and oligomeric assemblies, subsets of which could be detected with different antibody pairs in ultrasensitive immunoassays. However, to absolutely quantify mHTT, wildtype HTT or total HTT, many different antibody pairs and protein standards would have to be used and interpreting overlapping signals in these assays to absolutely define the precise mHTT proteoform composition in a sample would be very challenging. Another caveat in capturing different HTT proteoforms in immunoassays it that HTT antibody generation has been historically focused on targeting the N-terminal region of the protein, especially epitopes within the exon 1 region of the protein. It is possible that C-terminal fragments of HTT that arise from different proteolytic cleavage events are still not accounted for with the antibodies currently used in these assays. HTT is a protein scaffold in both its wildtype and disease forms, forming complex 3D structural assemblies of multi-protein complexes. Whether any of the interactions are maintained for extracellular HTT in different biofluid samples is unclear. This is an important consideration for HTT detection assays as some HTT protein-protein interactions almost certainly shield or occlude HTT antibody-epitope binding and thus alter assay signal. We demonstrate that buffer conditions of different stringency can alter assay signal arising from apo HTT compared to HTT in complex with HAP40. Ensuring all proteoforms are detected would be critical for absolute determination of “total” HTT protein levels, which cannot be achieved with current technology.

Our data demonstrate that mHTT detection assay signal is influenced by polyQ tract length, corroborating the findings of others (Vauleon et al., 2023). MW1 interaction with HTT is dependent on the polyQ tract length, showing preference but not specificity for disease-length polyQ tracts for both the denatured and native full-length HTT protein. This finding means that even measuring a single form or fragment of mHTT in a cohort of patient samples would be very difficult given the variation of polyQ tract length between individuals and even within a single patient sample due to variation over the disease course which arises due to somatic expansion (Aviolat et al., 2019). This mismatch in polyQ tract length protein standards and biological samples accounts for the mHTT detection paradox where mHTT levels exceed total HTT quantified in a single sample due to vastly different antibody-protein stoichiometry and therefore assay signal between antibody pairs used to detect HTT. This finding is applicable to other polyQ containing proteins where polyQ tract expansion also occurs during disease as we demonstrate with our analysis of wildtype and SCA3 representative ataxin-3 proteins.

Even with these caveats in mind, ultrasensitive HTT detection assays still have an important role to play in our evaluation of HTT as a biomarker of HD, and for assessment of target engagement of HTT-lowering therapeutics. We propose the recommendations laid out in **Box 1**, which advocate for relative quantitation of HTT to be reported by such assays rather than reporting HTT protein concentration, and that mHTT depletion assays are reconsidered as an approach to measure wildtype HTT. Relative reporting of assay data will also allow data interoperability and comparison across different clinical and preclinical studies.

## Materials and Methods

### Protein Construct Information

All protein expression constructs used in this study have been previously described (Denis et al., 2023; Harding et al., 2019) and are available through Addgene. These include full-length HTT with C-terminal FLAG-tag (Q23, Q25, Q30, Q36, Q42, Q52. Q54, Q60 and Q66), full-length HTT with N-terminal FLAG-tag (Q23, Q66), full-length HAP40, and full-length ataxin-3 (Q10, Q80). For full construct details and Addgene accession numbers, see **Supplementary Table 1**.

### Protein Expression and Purification

Full-length HTT proteins and HTT-HAP40 protein complexes were produced as previously described (Harding et al., 2021, 2019) and all plasmids are available through Addgene (“available plasmids from Harding et al. (2019) Journal of Biological Chemistry,” 2024). Briefly, Sf9 cells were infected with P3 recombinant baculovirus and grown until viability dropped to 80–85%, normally ∼72 h post-infection. For HTT-HAP40 complex production, a 1:1 ratio of HTT:HAP40 P3 recombinant baculovirus was used for infection. Cells were harvested, resuspended in 20 mM HEPES pH 7.4, 300 mM NaCl, 5% (v/v) glycerol supplemented with protease inhibitors and benzonase, then lysed with multiple freeze–thaw cycles and clarified by centrifugation. Proteins were purified by FLAG-affinity chromatography. All samples were purified with a final gel filtration step, using a Superose6 10/300 column in 20 mM HEPES pH 7.4, 300 mM NaCl, 1 mM TCEP, 2.5% (v/v) glycerol. Fractions of the peaks corresponding to the HTT monomer or HTT-HAP40 heterodimer were pooled, concentrated to 1 mg/mL, aliquoted and flash frozen prior to use in downstream experiments. Sample purity was assessed by SDS-PAGE.

N586 HTT Q68 and fusion HTT Q68 recombinant proteins were generated as previously described (Southwell et al., 2015). Briefly, the N586 fragment of the *HTT* gene was amplified by PCR of full-length HTT with BamHI and NotI restriction sites in the 5’ and 3’ primers, respectively. The PCR product was purified using the QiaQuick Gel Extraction kit (Qiagen) and digested with BamHI and Not1 to create sticky ends. the pGEX-6p-1 expression vector was digested using BamHI and NotI and gel purified. The insert and vector were ligated and transformed into DH5 *E. coli* and plated overnight on LB-agar plates containing 100 μ g/mL ampicillin. Colonies were screened by Qiagen mini-prep and confirmed by sequencing. The newly generated plasmids were then transformed into BL21 DE3 *E. coli* that were grown to OD 600 values of 0.8 and induced for protein production by IPTG. Cultures were lysed, and recombinant protein isolated by GST column purification and buffer exchanged into PBS using Amicon Ultra 10K MWCO centrifugal filters (Millipore). Recombinant proteins were quantified using a BCA assay (Pierce) and checked for purity using silver-stained SDS-PAGE gels.

Ataxin-3 proteins were produced as previously described (Denis et al., 2023). Ataxin-3 Q10 was overexpressed in E. coli BL21 CodonPlus (DE3) (Agilent). Ataxin-3 Q80 was produced by baculoviral transduction of this construct in Sf9 insect cell culture. For both proteins, harvested cell pellets were resuspended in 20 mM HEPES pH 7.4, 300 mM NaCl, 5% (v/v) glycerol, 1 mM TCEP supplemented with protease inhibitors and benzonase. The cell suspension was lysed by sonication and the clarified lysate was incubated with Talon resin (Cytiva). Resin was washed with a purification buffer supplemented with 5 mM imidazole and proteins eluted with a purification buffer supplemented with 300 mM imidazole. Eluted proteins were further purified by gel filtration using a S200 16/60 column equilibrated in the purification buffer. All samples were aliquoted, and flash frozen in liquid nitrogen prior to use. Protein purity was confirmed by SDS-PAGE.

### Mapping of HDB4 epitopes

Expression vectors to produce exon 1, N171, and N586 HTT proteins were generated by amplifying the indicated regions from full-length human HTT template DNA (Q68) by PCR, using primers with EcoRI and NotI restriction sites in the 5’ and 3’ primers respectively (**Supplementary Table 1**). The PCR products were purified by gel extraction using the QiaQuick gel extraction kit. The PCR products and pCI-Neo mammalian expression vector were then digested using EcoRI and NotI, gel purified, ligated, and transformed into Max Efficiency DH5α E. coli cells (Invitrogen #18258-012), then plated overnight on LB-agar plates containing 100 μg/mL ampicillin. Colonies picked from these plates were grown overnight in LB with 100 μg/mL ampicillin, then plasmid DNA was purified (Qiagen miniprep kit), and screened by restriction digest with EcoRI and NotI. Clones with expected sizes on restriction digest were confirmed by Sanger sequencing, then cultures were grown for large-scale DNA purification (Promega Maxiprep Kit). HEK293 cells were plated at 3×105 cells per well in a 6-well plate and grown overnight to 80% confluence, then transfected (Lipofectamine 2000). Alongside the Exon1, N171, and N586 constructs, pmaxGFP vector (Lonza) was included as a positive control for transfection, and strong GFP expression was observed 20 hours post-transfection. Cell pellets were harvested and lysed in SDP plus protease inhibitors. Proteins were quantified by DC assay, and SDS-PAGE was used for confirmation of protein purity and subsequent Western blotting with BKP1 and HDB4E10 antibodies.

### Immunoprecipitation-Flow Cytometry (IP-FCM)

The IP-FCM technique has been previously described (Schrum et al., 2007; Southwell et al., 2015). Briefly, capture antibodies were coupled to 5 μm CML latex microbeads (Invitrogen) and counted on a hemocytometer before storage at 4°C. Probe antibodies were biotinylated using EZ-Link Sulfo-NHS-Biotin (Thermo Scientific), free biotin removed by buffer exchange in Amicon Ultra 3K MWCO spin columns (Millipore), and antibody concentration brought to 0.5 mg/ml before storage at 4°C in PBS. Protein samples were diluted 1000 fM unless otherwise stated. CSF samples were diluted 1:1 to a total volume of 100 μl per replicate. Approximately 10^4^ beads in 5 μl NP-40 buffer (150mM NaCl, 50mM Tris (ph7.4), Halt Phosphatase and Protease inhibitors (10ul/ml), 0.5 M EDTA, 2mM Sodium Orthovanadate, 10mM NaF, 10mM Iodoacetamide, Surfact-Amps NP-40 1% (v/v) (Thermo Scietific, CAT#28324)) were mixed with 25 μl of recombinant protein in aCSF (125 mM NaCl, 2.5 mM KCL, 1.25 mM NaH2PO4, 1 mM MgCL2, 26 mM NaCO3, 2 mM CaCl2, 25 mM Dextrose) and incubated overnight at 4°C with rotation to prevent beads settling out of suspension. Beads were then washed in IP-FCM buffer (100 mM NaCl, 50 mM Tris pH 7.4, 1 % (w/v) bovine serum albumin (Sigma), 0.01 % (w/v) sodium azide) and incubated with biotinylated probe antibodies for 2 h, followed by another wash in IP-FCM buffer, incubation with 1:200 Streptavidin-PE (BD Biosciences) for 1 h, a final wash, and measurement on an Accuri flow cytometer (BD Biosciences). Bead doublets were gated out based on forward scatter area vs. forward scatter height plots, and a singlet bead gate was defined based on forward scatter height vs. side scatter height. All samples were run in three replicates, and the average of the median fluorescence intensity in the FL2 channel in the singlet bead gate indicated the abundance of HTT in the sample.

### Enzyme-Linked Immunosorbent Assay (ELISA) Analysis of HTT Allelic Series and Ataxin-3 proteins

ELISAs were conducted as previously described with some adaptations (Denis et al., 2023). Full-length HTT samples corresponding to an allelic series (Q-lengths 23, 25, 30, 36, 42, 52, 54, 60, 66) or Ataxin-3 samples (Q-lengths 10, 80) were quantified using the Pierce BCA Protein Assay kit (Thermo Scientific) as per manufacturer’s protocol. All proteins were diluted to 1 μg/mL using gel filtration buffer (20 mM HEPES pH 7.4, 300 mM NaCl, 2.5% glycerol, 1 mM TCEP at pH 7.4) and incubated in 96-well Nunc Maxisorp plates (Thermofisher Scientific, cat#442404) for 16 h at 4°C. Plates were washed four times with PBS with 0.005% (v/v) Tween-20 (PBS-T 0.005%) and blocked with PBS-T 0.005% with 1% (w/v) BSA (blocking buffer) for 2 h at 37°C and then for 3 h at 4°C. Plates were washed four times then incubated for 16 h at 4°C with 12-point 1:3 serial dilution of anti-polyQ MW1 (DSHB) in with each concentration in triplicate. The plate was then washed four times with blocking buffer and incubated for 1 h at 37°C with HRP-conjugated goat anti-mouse IgG (H+L) secondary antibody (1/50 000, Invitrogen, cat# 31430). After washing six times with blocking buffer, 100 μL of 1X TMB substrate (Invitrogen) was added per well and incubated at RT for ∼15 mins. The reaction was then quenched with 100 μL of 1 M phosphoric acid. The absorbances were measured at 450 nm using the BioTek Gen5 microplate reader (ThermoFisher Scientific). The following four negative control conditions were tested in triplicate wells to determine the total background signal: no HTT protein, no primary antibody, no secondary antibody, and washing buffer only. After defining specific binding as the absorbance values after subtracting the average absorbance of these control wells, the data was fitted to specific binding with hill slope using GraphPad Prism version 9.5.1.

### Western Blot Analysis of HTT Allelic Series

Full-length HTT samples corresponding to an allelic series (Q-lengths 23, 25, 30, 36, 42, 52, 54, 60, 66) were quantified using the Pierce BCA Protein Assay kit (Thermo Scientific) as per manufacturer’s protocol. 5 or 50 ng of each sample was loaded per lane on NuPAGE 4-12% Bis-Tris SDS-PAGE (Invitrogen) in 1X NuPAGE MOPS SDS running buffer (Invitrogen) for 3 h at 120 V. The proteins were then transferred onto 0.22 μM PVDF membranes (Bio-Rad) for 6 h at 30 V and 4°C. The membranes were blocked with 5% (w/v) milk powder in PBS with 0.1% (v/v) Tween-20 (PBS-T 0.1%) for 1 h at room temperature (RT), washed 3 times with PBS-T 0.1%, and then incubated with anti-polyQ MW1 (1/2000; DSHB) and anti-HTT EPR5526 (1/10,000; Abcam) for 16 h at 4°C with rocking. After three washing steps, the membrane was probed with secondary antibodies goat-anti-rabbit IgG-IR800 (1/3000, LI-COR) and donkey anti-mouse IgG-IR680 (1/3000, LI-COR) for 1 h at RT with rocking. The Odyssey CLx imaging system (LI-COR) was used to image the membrane and ImageStudio (LI-COR) was used for signal quantitation.

### MW1 depletion of Hu97/18 brain lysates and downstream analysis by western blot and MW1/HDB4 IP-FCM

Hu97/18 mice were killed with an overdose of intraperitoneal avertin and brains removed and placed on ice for ∼1 min to increase tissue rigidity. Olfactory bobs and cerebella were removed and the forebrain isolated, divided into hemispheres, and snap frozen in liquid nitrogen before storage at -80°C until use. Forebrain hemisphere were lysed by mechanical homogenization in NP-40 buffer, incubation on ice for 15 min., sonication at 25% for 5s, and removal of debris by centrifugation at 14,000xG for 10 min at 4°C. MW1 conjugated to magnetic dynabeads was incubated with 40 μg of total protein overnight at 4°C with gentle rotation. Beads were then isolated with a magnet and the flow through collected. The immunoprecipitation (beads) was separated on a low bis-acrylamide gel as previously described (Carroll et al., 2011) along with the flow through, and 40 μg of total lysate protein (input). Protein was transferred to nitrocellulose membrane, blocked for 1 hr at RT in 10% powdered milk in PBS, and probed HTT (MAB2166, Millipore) and calnexin (Sigma C4731) as a loading control. Primary antibodies were detected with IR dye 800CW goat anti-mouse (Rockland 610-131-007) and AlexaFluor 680 goat anti-rabbit (Molecular Probes A21076)-labelled secondary antibodies, and the LiCor Odyssey Infrared Imaging system. Band intensity was measured using densitometry and normalization to calnexin loading control.

## Acknowledgements

The Structural Genomics Consortium is a registered charity (no: 1097737) that receives funds from AbbVie, Bayer AG, Boehringer Ingelheim, Genentech, Genome Canada through Ontario Genomics Institute [OGI-196], the EU and EFPIA through the Innovative Medicines Initiative 2 Joint Undertaking [EUbOPEN grant 875510], Janssen, Merck KGaA (aka EMD in Canada and US), Pfizer, Takeda and the Wellcome Trust [106169/ZZ14/Z]. RJH receives funds from the Hereditary Disease Foundation and NSERC (RGPIN-2024-05769). ALS is supported with funds from NINDS (R01NS116099).

## Notes

### Competing Interest Statement

The authors have declared no competing interest.

