## Supplementary Information for "Challenges and advances for huntingtin detection in cerebrospinal fluid: in support of relative quantification"

***Supplementary Table 1. Primer sequences used in this study.***

| ***Primer*** | ***Sequence*** |
|---|---|
| Forward - HTT aa. 1 BamHI | GATCGGATCCATGGCGACCCTGGAAAAGCTG |
| Reverse - HTT aa. 586 NotI | GATCGCGGCCGCGTCTAACACAATTTCAGAACTGTC |
| Forward - pCI Neo CMV | GCTCACATGGCTCGACAGATCTTCA |
| Reverse - pCI Neo CMV | CTAGTTGTGGTTTGTCCAAACTCATC |
| Forward - HTT aa. 1744 EcoRI | GATCGAATTCAGGTTTCTATTACAACTGGTTG |
| Forward - HTT aa. 1744 EcoRI + ATG | GATCGAATTCATGAGGTTTCTATTACAACTGGTTG |
| Reverse - HTT aa. 2234 NotI | GATCGCGGCCGCGACCACCACCAGGTACTGTGC |
| Forward - HTT aa. 1 EcoRI | GATCGAATTCATGGCGACCCTGGAAAAGCTG |
| Reverse - HTT aa. 90 NotI | GATCGCGGCCGCGTCGGTGCAGCGGCTCCTC |
| Reverse - HTT aa. 171 NotI | GATCGCGGCCGCCTCGAGCTGTAACCTTGGAAG |

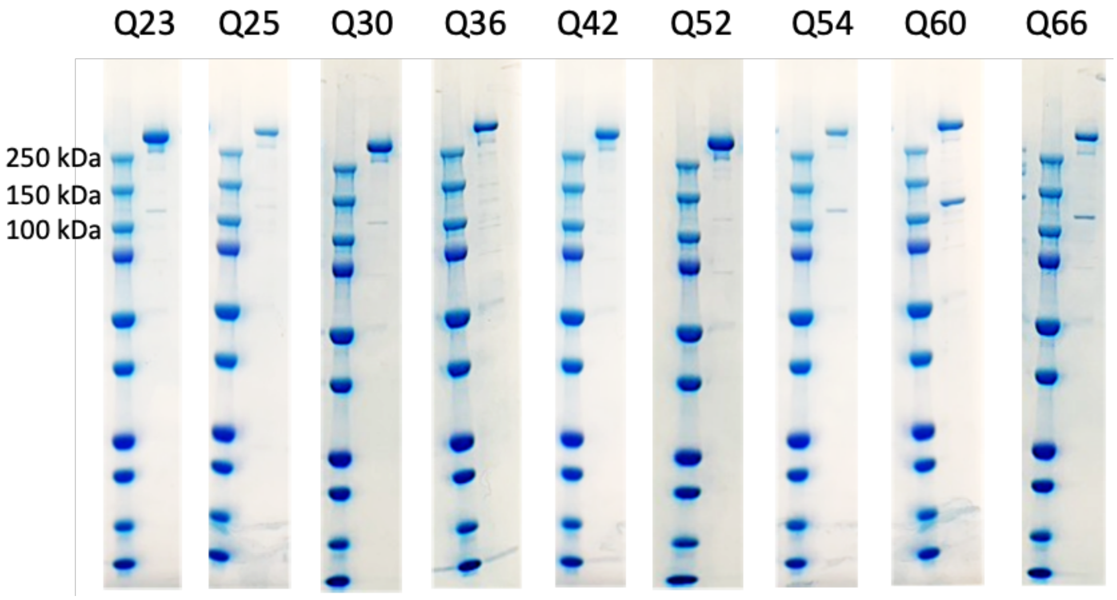


***Supplementary Figure 1. SDS-PAGE analysis of purified HTT allelic series. ~****2-5 µg of each HTT protein assessed by 4-20% Tris-Glycine SDS-PAGE showing >85% purity of all samples as determined by densitometry analysis.*

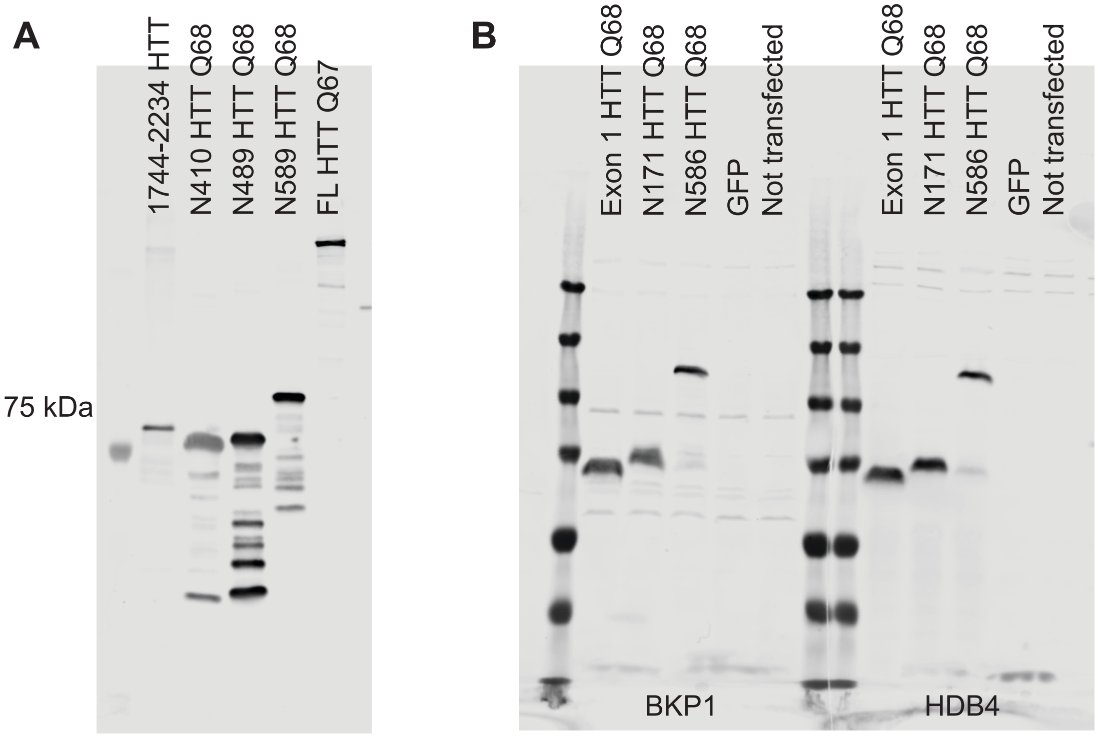


***Supplementary Figure 2. Mapping HDB4 epitopes with recombinant HTT proteins.*** *A. Recombinant HTT fragments, including a C-terminal fragment (aa. 1744-2234) that includes the immunogen used to raise HDB4 (aa. 1844-2131), N410, N489, and N589 as well as full-length HTT were used to investigate the specificity of HDB4 in denatured conditions. HDB4 recognizes full-length HTT and the C-terminal fragment as expected, but also the N-terminal fragments. B. Exon 1, N171, and N586 HTT constructs as well as a GFP construct were transfected into HEK293 cells. Cell lysates were separated by PAGE, transferred to nitrocellulose, and blotted with either BKP1 (left), which recognizes the first 17 N-terminal amino acids of HTT or HDB4 (right). We found that both antibodies recognized all three HTT fragments, but not GFP. Faint bands above the 250 kDa marker, likely the FL endogenous cellular HTT protein, were also seen in all lanes of both blots, including the not transfected cell lysate.*

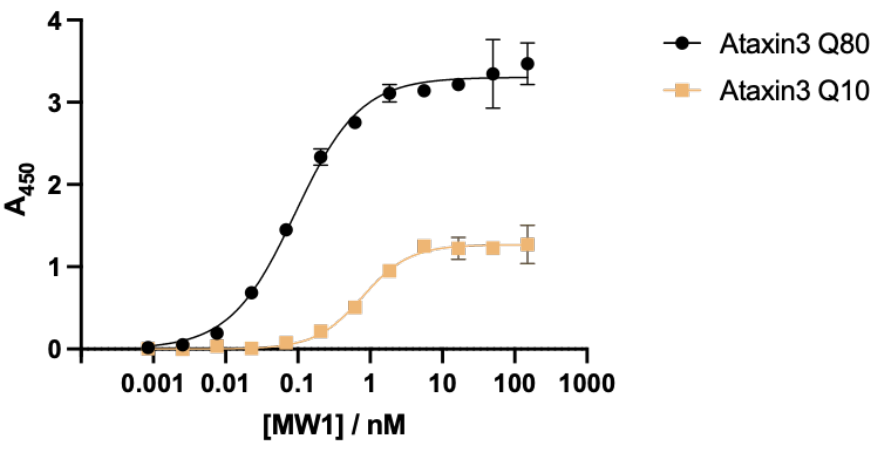

***Supplementary Figure 3. ELISA analysis of MW1-ataxin-3 interaction with different polyQ tract lengths.*** *Representative ELISA showing binding profile of MW1 to full-length Ataxin-3 with Q10 or Q80. Error bars are S.D. of three intra-assay replicates. Data fitted in GraphPad Prism with specific binding with hill slope model.*

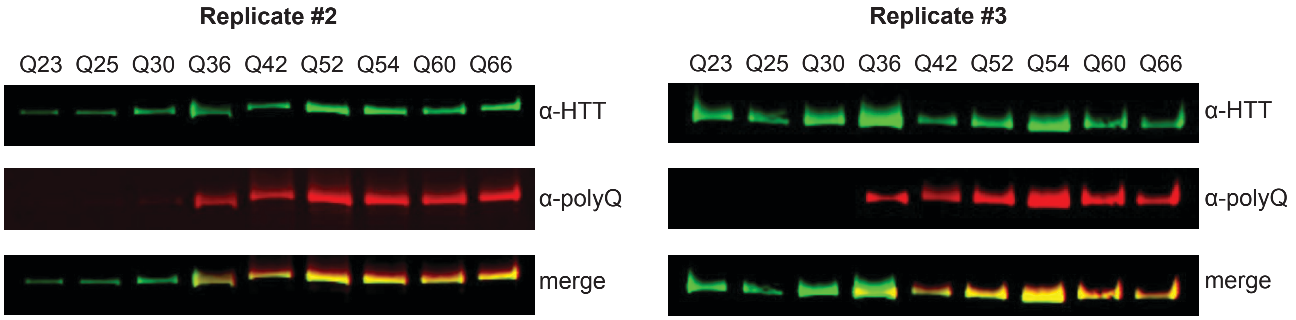


***Supplementary Figure 4. Complete western blot analysis of full-length HTT allelic series proteins with MW1 an EPR5526 antibodies.*** *Two replicates in addition to data shown in* ***Figure 2C*** *of western blot analysis of full-length HTT allelic series spanning Q23 to Q66 with ~5 ng loaded per lane. Blots probed with both α-HTT EPR5526 and α-polyQ MW1 shown separately and merged.*
